# Changes in insulin resistance do not occur in parallel with changes in mitochondrial content and function in male rats

**DOI:** 10.1101/2020.07.06.190702

**Authors:** Amanda J Genders, Jujiao Kuang, Evelyn C Marin, Nicholas J Saner, Javier Botella, Macsue Jacques, Glenn K McConell, Victor A Andrade-Souza, Javier Chagolla, David J Bishop

## Abstract

**Aims/hypothesis:** To investigate if there is a causal relationship between changes in insulin resistance and mitochondrial respiratory function and content in rats fed a high fat diet (HFD) with or without concurrent exercise training. We hypothesised that provision of a high fat diet (HFD) would increase insulin resistance and decrease mitochondrial characteristics (content and function), and that exercise training would improve both mitochondrial characteristics and insulin resistance in rats fed a HFD.

**Methods:** Male Wistar rats were given either a chow diet or a high fat diet (HFD) for 12 weeks. After 4 weeks of the dietary intervention, half of the rats in each group began eight weeks of interval training. *In vivo* glucose and insulin tolerance was assessed, as was *ex vivo* glucose uptake in epitrochlearis muscle. Mitochondrial respiratory function was assessed in permeabilised soleus and white gastrocnemius (WG) muscles. Mitochondrial content was determined by measurement of citrate synthase (CS) activity and protein expression of components of the electron transport system (ETS).

**Results:** HFD rats had impaired glucose and insulin tolerance. HFD did not change CS activity in the soleus; however, it did increase CS activity in WG (Chow 5.9 ± 0.5, HFD 7.2 ± 0.7 mol h^-1^ kg protein^-1^). Protein expression of components of the ETS and mitochondrial respiratory function (WG Chow 65.2 ± 8.4, HFD 88.6 ± 8.7 pmol O2 s^-1^ mg^-1^) were also increased by HFD. Exercise training improved glucose and insulin tolerance in the HFD rats. Exercise training did not alter CS activity in either muscle. Mitochondrial respiratory function was increased with exercise training in the chow fed animals in soleus muscle, but not in WG. This exercise effect was absent in the HFD animals. Mitochondrial characteristics did not consistently correlate with insulin or glucose tolerance.

**Conclusions/interpretation:** HFD induced insulin resistance, but it did not negatively affect any of the measured mitochondrial characteristics. Exercise training improved insulin resistance, but without changes in mitochondrial respiration and content. The lack of an association between mitochondrial characteristics and insulin resistance was reinforced by the absence of strong correlations between these measures. Our results suggest that defects in mitochondrial respiration and content are not responsible for insulin resistance in HFD rats.

## Introduction

Patients with insulin resistance or type 2 diabetes have been reported to have decreased skeletal muscle mitochondrial content [1–3], whilst patients with type 2 diabetes also display reductions in skeletal muscle mitochondrial respiratory function, when compared with healthy volunteers [2, 4–6]. However, despite these important cross-sectional observations, the relationship between skeletal muscle mitochondrial characteristics and insulin resistance continues to be debated and a causal relationship between altered mitochondrial characteristics and insulin resistance has yet to be established [7–9].

One approach to investigate the hypothesis that there is a causal relationship between altered mitochondrial characteristics and insulin resistance has been to induce insulin resistance in rodents and to determine whether there are also concomitant changes in mitochondrial characteristics. One popular model to induce insulin resistance is the provision of a high-fat diet (HFD), which causes whole-body insulin resistance after one week and skeletal muscle insulin resistance after three weeks [10]. However, the studies conducted to date have generated conflicting results, with both decreased [11–13] and increased mitochondrial respiratory function [14, 15] reported following a HFD. A potential factor contributing to these contrasting findings is that many of the studies have assessed mitochondrial respiratory function in isolated mitochondria [13, 14]. Despite the wide use of this technique, standard methods do not isolate the entire mitochondrial pool and this may result in a biased representation of the mitochondrial pool [16]. The isolation protocol also removes mitochondria from their cellular environment, fragments mitochondria, and can alter respiration and hydrogen peroxide production [16]. More research is required to investigate whether interventions that cause insulin resistance (e.g., HFD) also decrease mitochondrial respiration, using techniques that assess mitochondrial respiratory function *in situ*.

If sub-optimal mitochondrial characteristics cause insulin resistance, then interventions that improve mitochondrial characteristics (e.g., increase markers of mitochondrial content and/or respiratory function) should improve insulin resistance. We and others have reported that exercise training is a powerful stimulus that increases both mitochondrial content and respiratory function [17–22]. However, while a small number of studies have examined the ability of exercise to prevent or reverse various aspects of the metabolic syndrome induced by HFD [23–25], we are not aware of any studies that have investigated whether mitochondrial characteristics affected by HFD are reversed by exercise training. Such a study would help clarify the relationship between altered mitochondrial characteristics and insulin resistance, but would also have implications for understanding whether the benefits of exercise training in a model of lipid oversupply (i.e., HFD) can be attributed to changes in mitochondrial characteristics.

Therefore, the aim of this study was to investigate the effects of 12 weeks of a HFD followed by eight weeks of interval training on both insulin resistance and mitochondrial characteristics (respiratory function measured *in situ* and markers of mitochondrial content). If changes in insulin resistance occur in parallel with changes in mitochondrial content and/or mitochondrial respiratory function, this would provide support for a causal relationship between altered mitochondrial characteristics and insulin resistance. We hypothesised that male rats fed a HFD would develop insulin resistance, as assessed by glucose and insulin tolerance tests and *ex vivo* glucose uptake, and this would be accompanied by decreased mitochondrial content and respiratory function. We further hypothesised that exercise training would improve both mitochondrial respiratory function and insulin resistance in rats fed a HFD.

## Methods

### Animals and experimental overview

The Victoria University Animal Ethics Committee approved this study (AEC 15/002). All procedures were performed according to the Australian Code of Practice for the Care and Use of Animals for Scientific Purposes (National Health and Medical Research Council, Australia, 8^th^ Edition). Male Wistar rats were purchased from the Animal Resource Centre (Perth, WA) at four weeks of age. Rats were housed in pairs in a temperature-controlled room and maintained with a chow diet (Specialty Feeds, Perth, WA) and drink *ad libitum* on a 12:12 h light-dark cycle, 18-22C, with approximately 50% humidity. The rats were provided with environmental enrichment including shredded paper, cardboard, and plastic pipes. The intervention began at six weeks of age (216 ± 22 g), with rats randomly assigned to each group. Eighteen rats were maintained on the regular chow diet, while 18 rats were given a 43% high-fat diet (SF04-001, Specialty Feeds, WA) verified independently by FeedTest (Werribee, VIC). Not all animals were studied at the same time; however, rats from all four groups were studied concurrently. The number of animals used in each group was determined from the expected variability of mitochondrial respiratory function, and previous experiments. At 19 weeks of age (at least 72 h after the completion of the last exercise bout) rats were killed using 90 mg.kg^-1^ intraperitoneal (IP) pentobarbitone (the ethically approved method at our university) and the muscles quickly removed for analysis (epitrochlearis, soleus, white portion of the medial gastrocnemius). Freshly dissected muscle was used for mitochondrial respiration and *ex vivo* glucose uptake analysis, with the remainder snap-frozen in liquid nitrogen and stored at −80 °C for further analysis (Fig. 1). Most human studies measure changes in the one muscle, the vastus lateralis, while the majority of studies in mice (due to their small size) investigating mitochondrial characteristics tend to use only one muscle, usually either the quadriceps or red gastrocnemius. This doesn’t take into account the influence of fibre type on measurements, which may vary between fibre types [26–29]. Thus, using a rat model in this study has allowed us to more fully explore mitochondrial characteristics and possible variations between different muscles. All exercise and other procedures took place in the daytime and in a separate laboratory space from where the animals were housed. Animals were carefully monitored after the IPITT and IPGTT procedures to ensure that entries made to obtain blood samples were not bleeding. The body mass and food intake of the rats was monitored throughout the experimental period.

**Figure 1:**
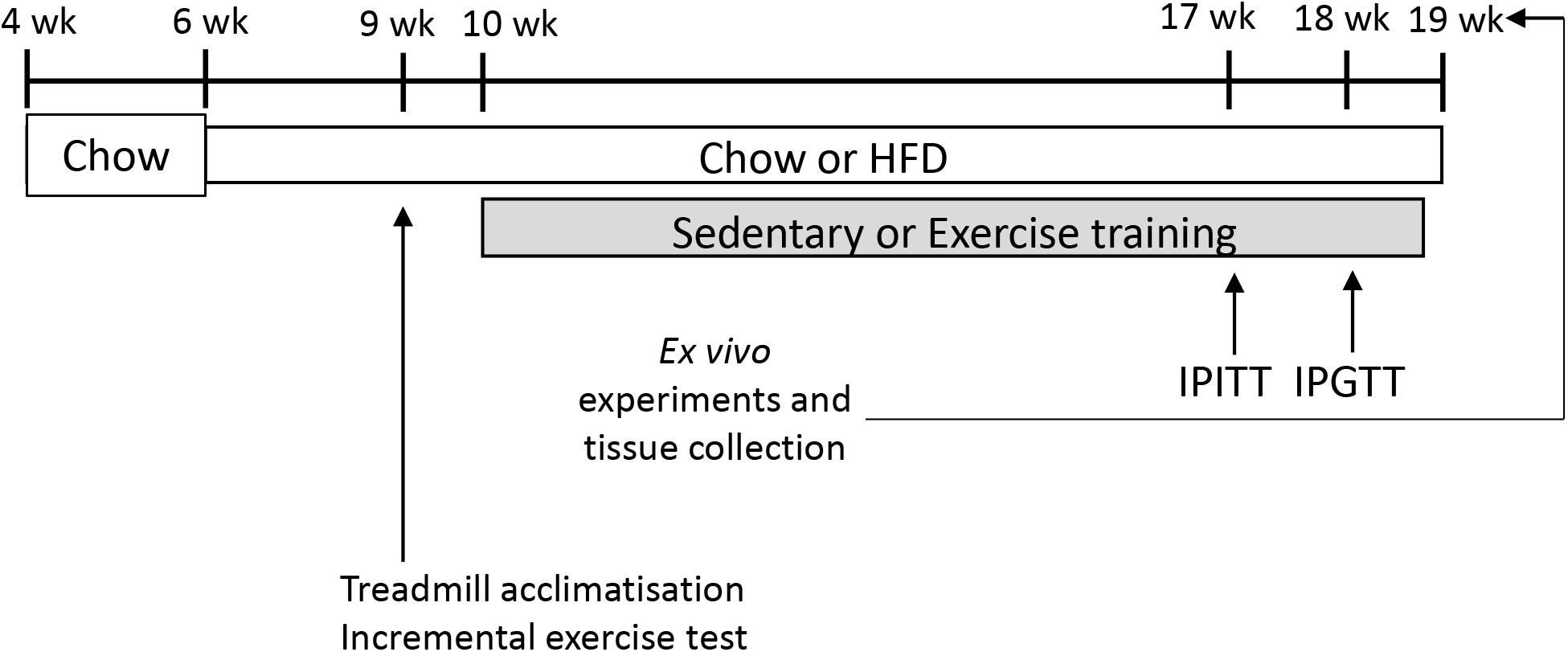
Outline of study design. Male Wistar rats were purchased at four weeks old and had two weeks acclimatisation prior to starting in the study at 6 weeks old, either remaining on a chow diet (n = 18) or changing to a high fat diet (HFD) (n = 18). At nine weeks of age, animals were acclimatised to the treadmill and undertook the incremental exercise test. After four weeks of the dietary intervention half the rats from each group started interval exercise training five times per week for eight weeks. At 17 weeks old all rats had an intraperitoneal insulin tolerance test (IPITT) and at 18 weeks old an intraperitoneal glucose tolerance test (IPGTT).

### Treadmill exercise

At nine weeks of age, all rats were acclimatised to the treadmill (Columbia Instruments, OH) over five separate sessions starting from a stationary treadmill at a five-degree incline to 15 m.min^-1^ at a 10-degree incline. Their exercise capacity was then assessed using an incremental exercise test. The incline of the treadmill was set at 10 degrees and the test was started at a speed of 9.6 m.min^-1^. The speed of the treadmill was increased 3 m.min^-1^ every three minutes. Animals were removed from the treadmill when they could no longer keep up with the speed despite encouragement from an air puff and/or touch. Exercise training began at 10 weeks of age and consisted of seven (1^st^ week) to 12 (weeks 6-8) 2-min intervals (interspersed with 1 min of rest) performed at 80% of the top speed reached in the incremental exercise test five days per week for the animals included in the exercise training groups. Intervals were undertaken at approximately 25 m.min^-1^ in the chow fed animals and approximately 23 m.min^-1^ in the HFD animals. Sedentary control animals were removed from their home cage and room at the same time as the exercising animals each day.

### Insulin tolerance test

An IPITT was performed in all animals at 17 weeks of age, after a 2 h fast. Insulin (0.75 IU.kg^-1^) was given IP and blood samples were collected immediately from the tail vein prior to the insulin dose and then 20, 40 and 60 minutes after injection. Blood glucose concentration was measured using a hand-held glucose monitor (Accu-Chek Performa Nano, Roche Diagnostics, Mannheim, Germany). n = 9 in all groups except the Chow Ex. group where for one animal the rat was not injected successfully with insulin.

### Glucose tolerance test

An IPGTT was performed in all animals at 18 weeks of age after a 5-6 h fast. Glucose (1g.kg^-1^) was given IP and blood samples were collected immediately from the tail vein prior to the glucose dose and then 15, 30, 45, 60 and 90 minutes after injection. Blood glucose concentration was measured using a hand-held glucose monitor (Accu-Chek Performa Nano, Roche Diagnostics, Mannheim, Germany). n = 9 in the Chow Ex. group, n = 8 in Chow Sed., HFD Sed., HFD Ex. groups due to unsuccessful injection of the glucose. Plasma insulin was measured using an ELISA, CV% = 6.73 (80-INSTRU-E01, Alpco, Salem, NH, USA) [30]. Insulin measurements where for n = 5-7 for each group due to missed glucose injection or insufficient blood sample at individual time points.

### Ex vivo glucose uptake

Ex vivo glucose uptake was measured in the epitrochlearis muscle as previously described [31] with the exception that 60 nmol.L^-1^ insulin was used. Some values were excluded from the analysis due to technical errors, therefore final analysis was conducted on n = 7-8 per group.

### Western blotting

Soleus and white gastrocnemius muscle samples were homogenised in ice cold lysis buffer (0.05 M Tris pH 7.5, 1 mM EDTA, 2 mM EGTA, 10% vol/vol glycerol, 1% vol/vol Triton X-100, 1 mM DTT) with a Protease and Phosphatase Inhibitor cocktail (Cell Signaling Technology, Danvers, MA). Proteins (5 to 10 μg per sample) were separated by electrophoresis (12% TGX Stain-Free FastCast Acrylamide gels) and then transferred onto PVDF membranes. Membranes were then blocked and then probed with an OXPHOS antibody cocktail (abcam 110413) primary antibody overnight at 4 °C at 1:1000 in TBST with 5% wt/vol BSA. Blots were then washed with TBST prior to incubation with HRP-linked anti-mouse secondary antibody (Perkin Elmer) for 1 h at room temperature. Blots were developed using Clarity ECL and visualised using a ChemiDoc™ MP (Bio-Rad Laboratories, Gladesville, Australia). All bands were quantified using ImageLab software (Bio-Rad Laboratories). Individual subunit protein expression was normalised to total protein. As used by Dirks et al [32] we also reported total ETS subunit protein content.

### Citrate synthase activity assay

Citrate synthase (CS) activity was used as a marker of mitochondrial content [33] and determined as described previously [17].

### Muscle fibre mitochondrial respiration

Researchers conducting these measures were blinded to experimental group. Muscle fibres were prepared as previously described [17, 34]. For all animals, the respiratory parameters of the total mitochondrial population of the soleus and white gastrocnemius muscle fibres (2-3 mg) were studied *in situ* using two separate high resolution Oxygraph-2k instruments (Oroboros Instruments, Austria) as previously described [17]. A substrate-uncoupler-inhibitor titration (SUIT) protocol was used [34], with the substrates and inhibitors added to the chamber as follows: 2 mM malate and 0.2 mM octanyolcarnitine were added in the absence of ADP for the measurement of leak respiration (ETF)_L_ through Complex I; 3 mM MgCl2 and 5 mM ADP were then added for the measurement of (ETF)_P_; 5 mM pyruvate addition allowed the measurement of Complex I and ETF_P_ respiration (maximal oxidative phosphorylation (oxphos) capacity (P)). This was followed by 10 mM succinate addition which allowed for the measurement of P through CI and II (CI + II)_P_ (measurement CV soleus muscle = 19.05%, TEM 29.62 pmol O2 s^-1^ mg^-1^; gastrocnemius muscle 15.82%, TEM 16.44 pmol O2 s^-1^ mg^-1^). Cytochrome c (10 μM) was then added to check for outer membrane integrity. Electron transport system (ETS) capacity through CI and II was determined by a number of titrations of carbonyl cyanide 4-(trifluoromethoxy)phenylhydrazone (FCCP) (0.5-1.5 μM) (CI+II_E_). Finally, Antimycin A (3.75 μM) was used to inhibit Complex III to allow for the measurement and correction of residual oxygen consumption, which is representative of non-mitochondrial oxygen consumption.

### Beta(3)-hydroxyacyl-CoA dehydrogenase activity assay

Beta(3)-hydroxyacyl-CoA dehydrogenase (β-HAD) activity was determined in triplicate in a 96-well plate at 30°C. 25 μL muscle lysate/lysis buffer, 5 μL 10% vol/vol Triton x-100 (diluted in 40% ethanol), and 215 μL assay mix (50 mM Tris pH 7.0, 2 mM EDTA, 0.375 mM NADH) was added to each well. The plate was then shaken in the x-Mark Microplate spectrophotometer (Bio-Rad) for 30 s. 5 μL 5 mM acetoacetyl-CoA was then added to each well. The plate was immediately returned to the plate reader where it was mixed again for 30 s, and absorbance was measured at 340 nm at intervals of 10 s for soleus muscle lysate, and 15 s for gastrocnemius muscle lysate for 50 reads. β-HAD activity was calculated and reported in mmol kg protein^-1^ h^-1^.

### Statistical analysis

Results in the figures are expressed as mean ± SD. Data were analysed for statistical significance using SPSS Statistics 22 using a two-way ANOVA for the main effects of diet or exercise training, and one-way ANOVA as appropriate with post hoc testing for differences between individual groups, with p ≤ 0.05 treated as significant. Data was checked for normal distribution and variance. In addition, basal vs insulin stimulated *ex vivo* glucose uptake was determined using a paired t-test for each treatment group. Effect sizes are reported as Cohen’s d. Mean ± 95% confidence intervals are reported for significant main effects in the results section text. Correlations were determined using a Pearson correlation.

## Results

### Body composition

Prior to the intervention, the rats from each group (± SD) had the following body mass: Chow Sed. 226 ± 23 g; Chow Ex. 221 ± 20 g; HFD Sed. 207 ± 23 g; HFD Ex. 210 ± 22 g. The HFD significantly increased total body mass (Chow 520 ± 26 g, HFD 570 ± 26 g, p = 0.003, ES = 1.1), whilst exercise training decreased body mass (Sed. 563 ± 26 g, Ex. 530 ± 26 g, p = 0.04, ES = 0.7) (Figure 2A). The HFD increased epididymal fat mass (Chow 6.16 ± 1.88 g, HFD 15.17 ± 1.99 g, p = 0.0001, ES = 2.5) (Figure 2B), but there was no significant effect of exercise training on epididymal fat mass.

**Figure 2:**
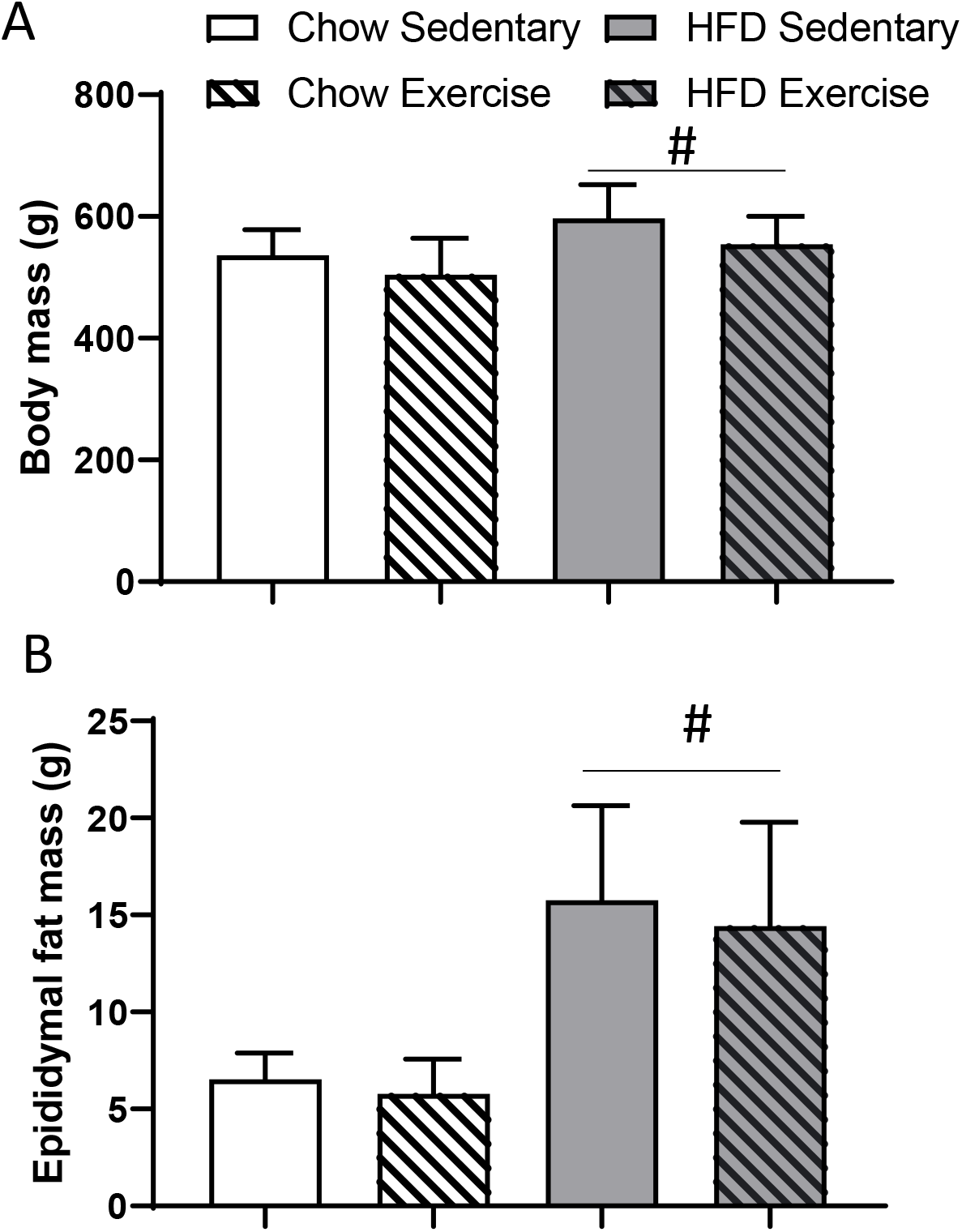
Body mass and epididymal fat pad mass. A. Body mass B. Epididymal fat mass. Mean ± SD, n = 9 for each group, #p≤ 0.05 main effect of HFD. There was a significant main effect of exercise training on body mass.

### Insulin and glucose tolerance

The HFD significantly decreased insulin tolerance (Chow 314 ± 17 mM*min, HFD 341 ± 16 mM*min, p = 0.031, ES = 0.8) (Fig 3A, B). Exercise training did not increase insulin tolerance in the chow-fed animals, but normalised insulin tolerance in the HFD animals (Fig. 3A,B). The HFD also significantly decreased glucose tolerance, as measured by an IPGTT (Chow 345 ± 67 mM*min, HFD 589 ± 66 mM*min, p = 0.0001, ES = 1.96). Glucose tolerance was not improved with exercise training (Fig. 3C,D); however, the increase in the insulin response to a glucose injection in the sedentary HFD animals was absent in the HFD animals who exercised (Chow Sed. vs HFD Sed. p = 0.039, ES = 1.6, Chow Sed. vs HFD Ex. p = 0.642, ES = 0.4) (Fig. 3E,F). *Ex vivo* glucose uptake was increased by insulin in each group, but was not altered by exercise training or HFD in the epitrochlearis muscle (Fig. 3G).

**Figure 3:**
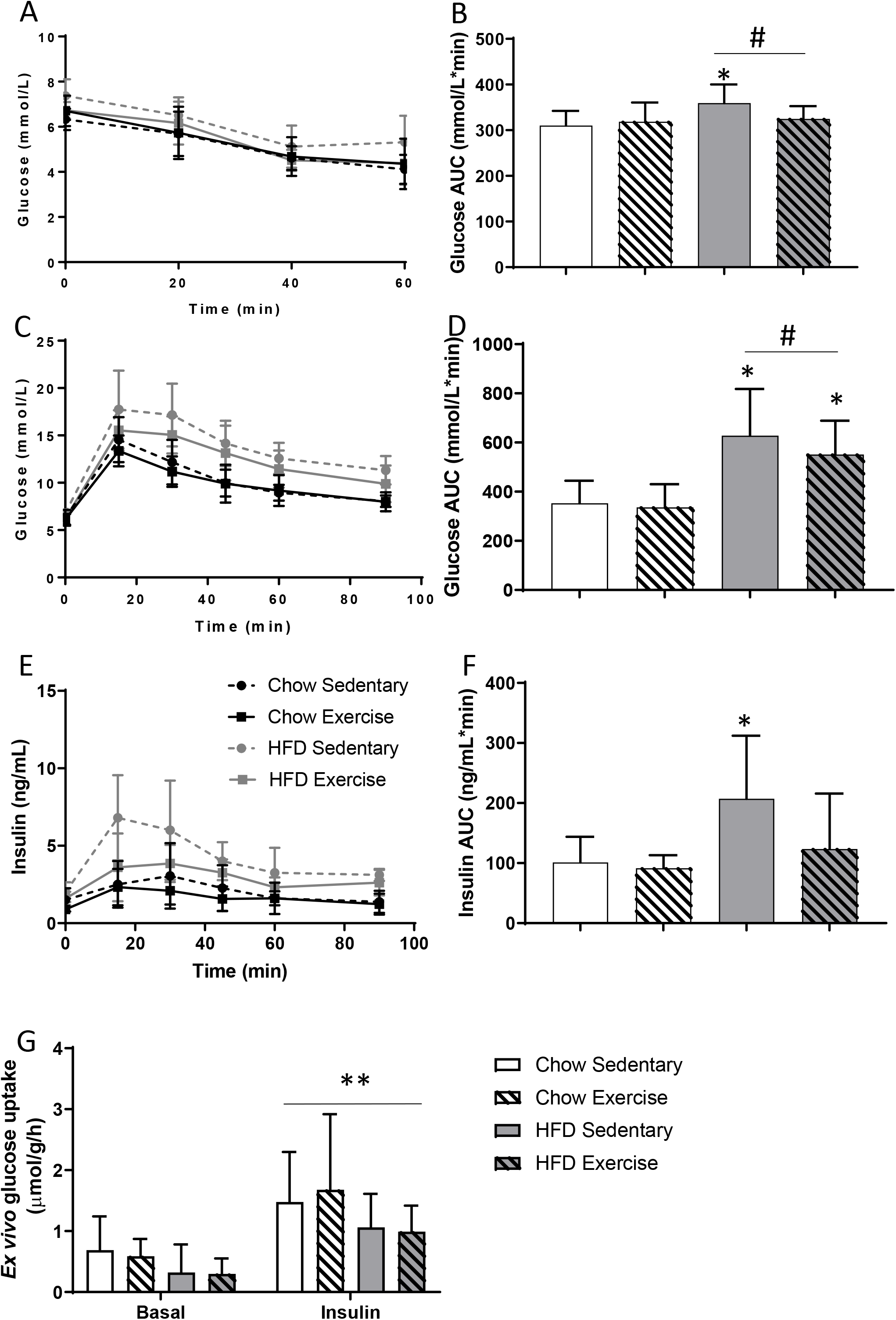
Insulin sensitivity and glucose tolerance is impaired in HFD rats. A. Blood glucose following intraperitoneal insulin injection – insulin tolerance test (IPITT) B. Glucose area under curve (AUC) for IPITT C. Blood glucose following i.p. glucose injection – glucose tolerance test (IPGTT) D. Glucose AUC for IPGTT E. Plasma insulin following i.p. glucose injection (IPGTT) F. Insulin AUC for IPGTT G. *Ex vivo* glucose uptake in epitrochlearis muscle. Mean ± SD, n = 9 for each group except for missing values described in the methods, *p≤ 0.05 vs chow sedentary group, # main effect of HFD. **p≤ 0.05 insulin increases glucose uptake compared to basal condition.

### Mitochondrial content

Citrate synthase (CS) activity was not altered in the soleus muscle by either exercise training or diet (Fig. 4A). However, the protein content of Complex I (Chow 1.1 ± 0.1, HFD 1.3 ± 0.16, p = 0.005, ES = 1.0) and III (Chow 1.2 ± 0.4, HFD 1.9 ± 0.4, p = 0.004, ES = 1.1) was greater in the soleus muscle after 12 weeks of HFD (Fig. 4C). This was reflected in the total OXPHOS protein content, which was greater after 12 weeks of HFD (Chow 1.05 ± 0.18, HFD 1.57 ± 0.21, p = 0.001, ES = 1.24) (Fig. 4C). Following the HFD, CS activity was greater in the gastrocnemius muscle (Chow 5.9 ± 0.5 mol/h/kg protein, HFD 7.2 ± 0.7 mol/h/kg protein, p = 0.007, ES = 1.0) (Fig. 4B), as was protein expression of components of the respiratory chain (CI Chow 1.0 ± 0.2, HFD 1.5 ± 0.2, p = 0.009, ES = 1.0) (CIII Chow 0.7 ± 0.2, HFD 1.73 ± 0.5, p = 0.0001, ES = 1.7) (Fig. 4D). This was also reflected in the total OXPHOS protein content, which was greater after 12 weeks of HFD (Chow 0.96 ± 0.14, HFD 1.47 ± 0.25, p = 0.002, ES = 1.18) (Fig. 4D). Exercise training did not alter CS activity or protein content of components of the respiratory chain in the gastrocnemius muscle of chow-fed rats (all p>0.05).

**Figure 4:**
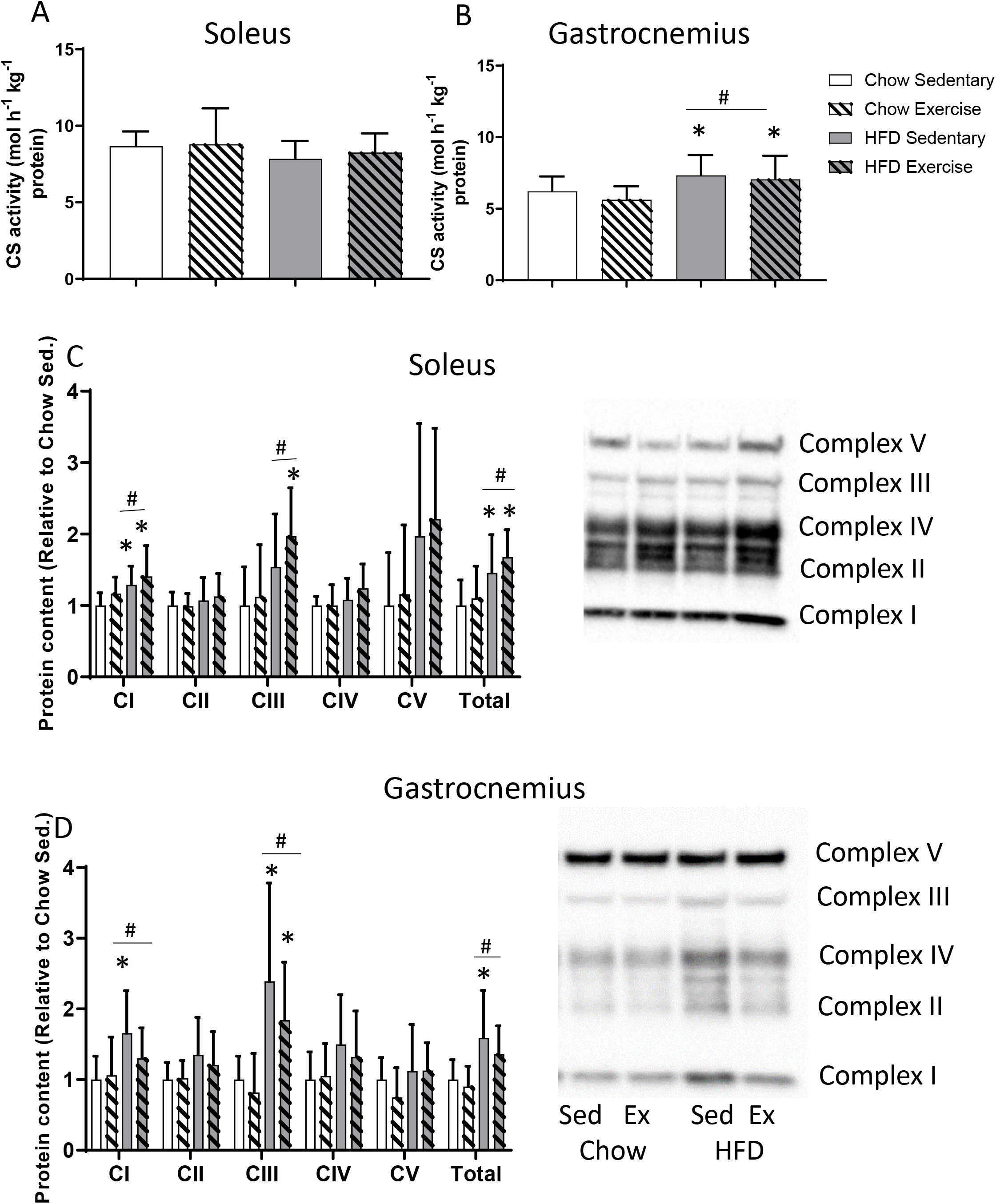
Citrate synthase (CS) activity and electron transport system (ETS) subunit protein content. A. CS activity in the soleus muscle B. CS activity in the white gastrocnemius muscle C. Protein content of subunits of ETS in the soleus muscle D. Protein content of subunits of ETS in the white gastrocnemius muscle (n = 7 Chow Ex, samples excluded to technical error). Mean ± SD, n = 9 for each group, *p≤0.05 vs chow sedentary group.

### Mitochondrial respiratory function

The HFD increased mitochondrial respiratory function (oxidative phosphorylation capacity through complex I and II; CI+II_P_) in both the soleus (Chow Sed. vs HFD Sed. p = 0.02, ES = 1.3, Chow Sed. vs HFD Ex. p = 0.037, ES = 1.2, no main effect of diet p = 0.111, ES = 0.6; Fig.5A) and the gastrocnemius muscles (Chow Sed. Vs HFD Sed. p = 0.006, ES = 1.3; Chow Sed. vs HFD Ex. p = 0.08, ES = 1.1; Chow 65.2 ± 8.4, HFD 88.6 ± 8.7 pmol O2 s^-1^mg^-1^, main effect of diet p = 0.0001, ES = 1.38; Fig.5B). There was no further increase in mitochondrial respiratory function in either muscle of the HFD + exercise group, but mitochondrial respiratory function was increased with exercise training in the chow fed animals in the soleus muscle (p = 0.045, ES = 1.1) (Fig. 5A) but not the gastrocnemius muscle (p = 0.408, ES = 0.6) (Fig. 5B).

**Figure 5:**
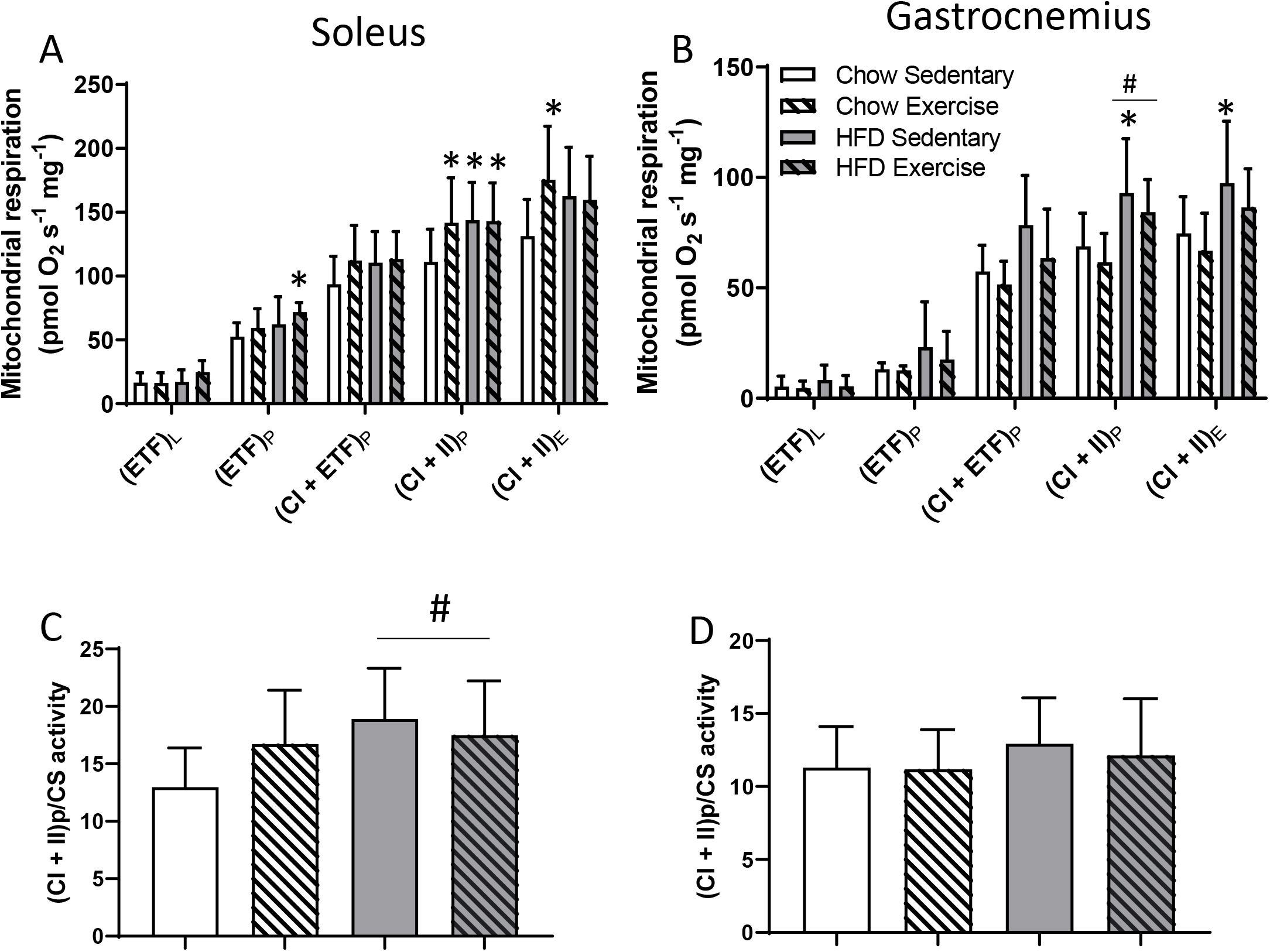
Mitochondrial respiratory function in response to a high fat diet (HFD) and exercise training. A. Mitochondrial respiratory function in the soleus muscle. B. Mitochondrial respiratory function in the white gastrocnemius muscle. C. Mitochondrial respiratory function in the soleus muscle normalised to citrate synthase (CS) activity. D. Mitochondrial respiratory function in the gastrocnemius muscle normalised to CS activity. CI: complex I; CI+II: convergent electron input through CI and CII; E: electron transport system (ETS) capacity; L: leak respiration; P: oxidative phosphorylation capacity Mean ± SD, n = 9 for each group with the exception of one muscle measurement for HFD Ex group (soleus and gastrocnemius), and one soleus measurement in the Chow Ex group due to non-responsive muscle, *p≤ 0.05 vs chow sedentary group. #p≤ 0.05 main effect of HFD.

When CI+IIP respiration was normalised to CS activity there was a significant effect of the HFD to increase CI+II_P_ respiration in the soleus muscle (Chow 14.7 ± 2.2, HFD 17.9 ± 2.1, p = 0.044, ES = 0.3) (Fig. 5C). There was no significant effect of the HFD on CI+II_P_ respiration when normalised to CS activity in the gastrocnemius muscle (Chow 11.23 ± 1.51, HFD 12.53 ± 1.56, p = 0.232, ES = 0.4) (Fig. 5D).

### B-HAD activity

To examine skeletal muscle mitochondrial fatty acid oxidative capacity, we measured the activity of β-HAD. β-HAD activity was significantly increased by the HFD in the soleus muscle (Chow 1840.3 ± 1097.3, HFD 4338.1 ± 1097.3 mol h^-1^kg^-1^ protein, p= 0.0001, ES = 1.2) (Fig. 6A), but was not altered by exercise training. β-HAD activity was not altered by the HFD or exercise training in the gastrocnemius muscle (Fig. 6B).

**Figure 6:**
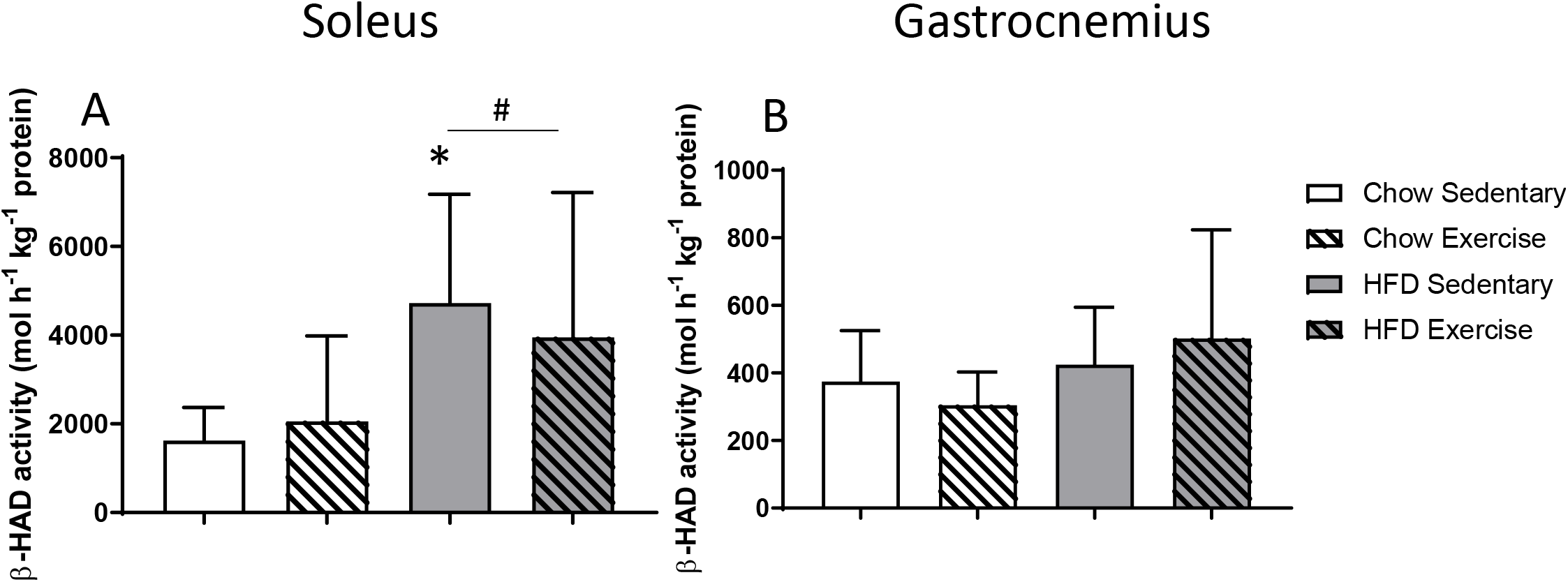
Beta-HAD activity in. A. the Soleus muscle, and B. White gastrocnemius muscle. Mean ± SD, n = 9 for each group, *p≤ 0.05 vs chow sedentary group.

### Correlations

The ITT AUC was positively correlated with CI + II oxidative phosphorylation (Fig. 7A), but not CS activity (Fig. 7E) in the soleus. In the gastrocnemius muscle, ITT AUC was correlated with CS activity (Fig. 7F) but not CI + II oxidative phosphorylation (Fig. 7B). The GTT AUC was not correlated with CI + II oxidative phosphorylation (Fig. 7C) or CS activity in the soleus (Fig. 7G). In the gastrocnemius the GTT AUC was correlated with CI + II oxidative phosphorylation (Fig. 7D), but not with CS activity (Fig. 7H). β-HAD activity in the soleus muscle was correlated with both AUC for GTT and ITT (Fig. 8A, B). β-HAD activity in the gastrocnemius muscle was correlated with AUC for the ITT, but not the GTT (Fig. 8C, D). Epididymal fat mass was not correlated with ITT AUC (Fig. 9A). However, epididymal fat mass was correlated with GTT AUC (Fig. 9B). Body mass was correlated with both ITT AUC (Fig. 9C) and GTT AUC (Fig. 9D).

**Figure 7:**
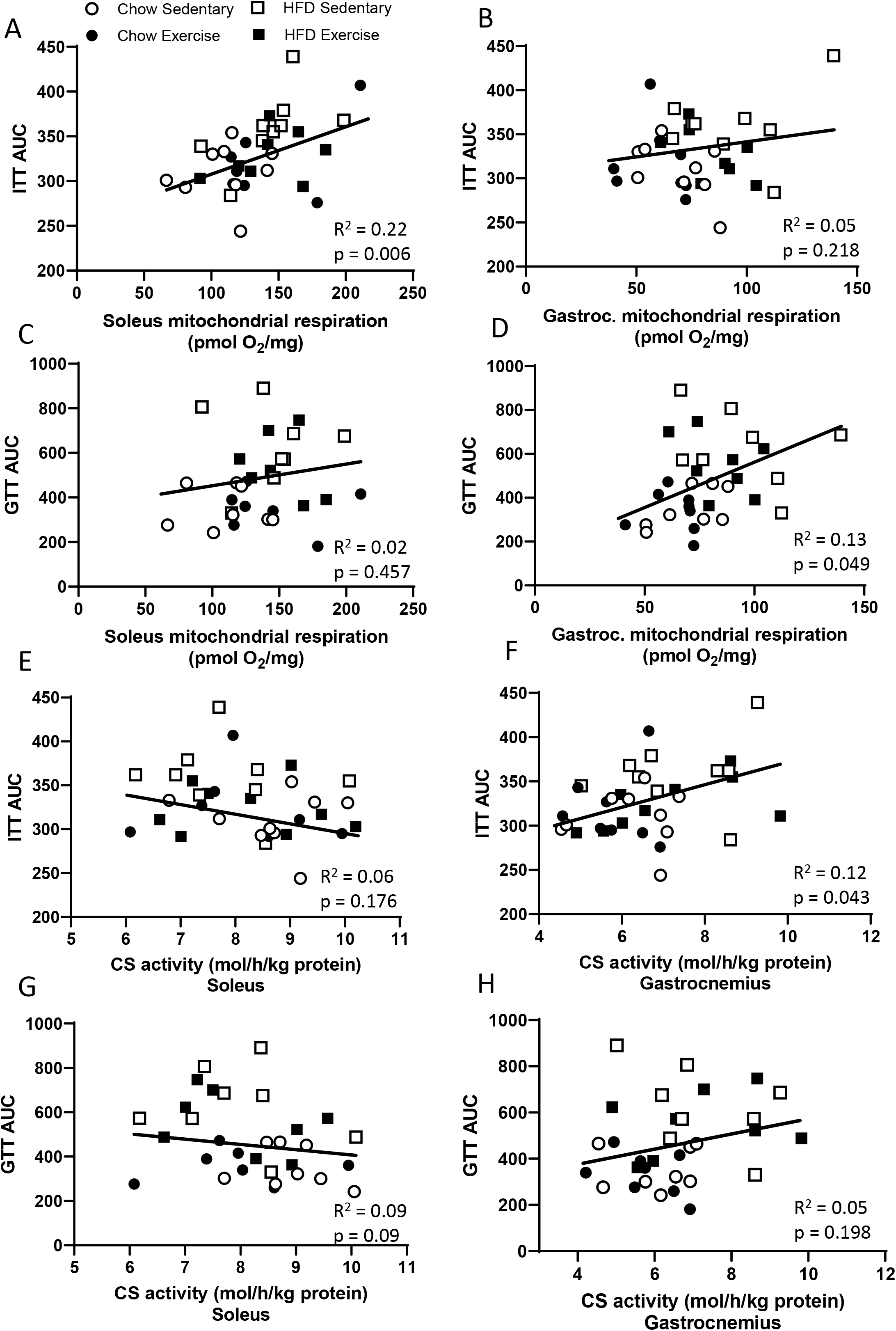
Correlations between metabolism and mitochondrial parameters. A. ITT glucose AUC and soleus (CI+II)p respiration. B. ITT glucose AUC and gastrocnemius (CI+II)p respiration. C. GTT glucose AUC and soleus (CI+II)p respiration. D. GTT glucose AUC and gastrocnemius (CI+II)p respiration. E. ITT glucose AUC and soleus CS activity (one soleus CS activity value was removed due to it being much higher than the other values for this correlation and with GTT AUC). F. ITT glucose AUC and gastrocnemius CS activity. G. GTT glucose AUC and soleus CS activity. H. GTT glucose AUC and gastrocnemius CS activity. n = 36.

**Figure 8:**
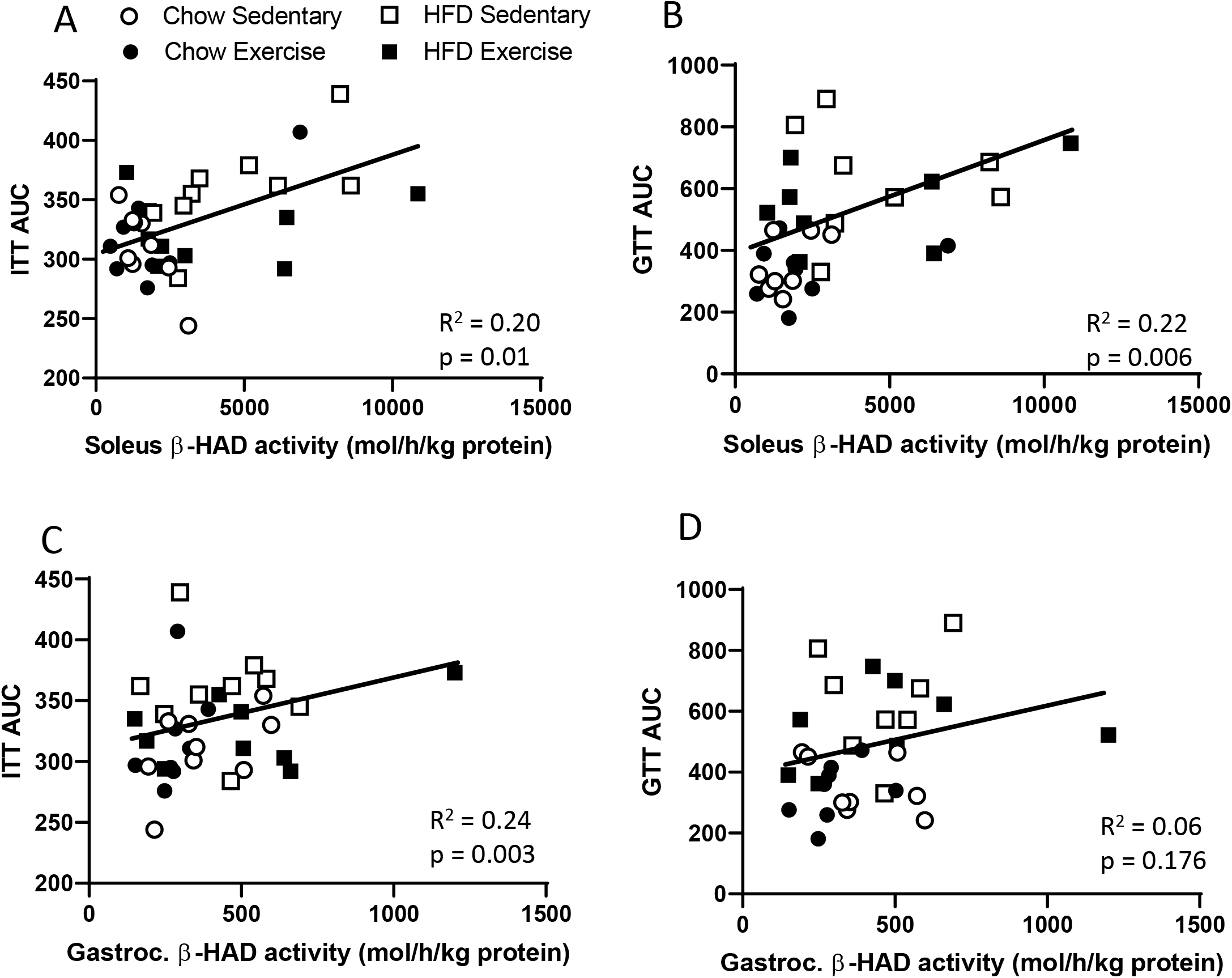
Correlations between metabolism and β-HAD activity. A. Soleus muscle β-HAD activity and ITT glucose AUC. B. Soleus muscle β-HAD activity and GTT glucose AUC. C. Gastrocnemius muscle β-HAD activity and ITT glucose AUC. D. Gastrocnemius muscle β-HAD activity and GTT glucose AUC. n = 36.

**Figure 9:**
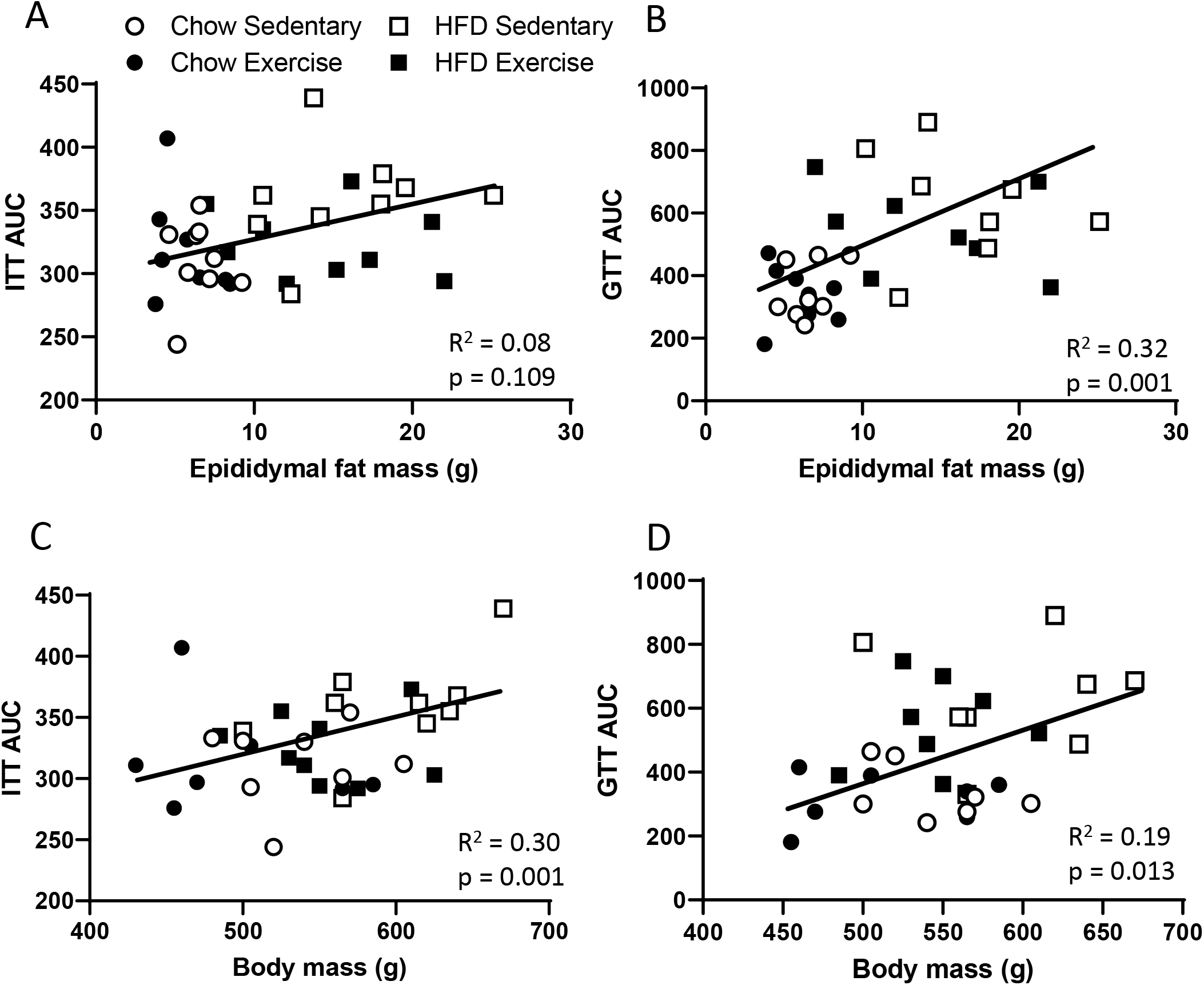
Correlations between metabolism and fat and body mass. A. Epididymal fat mass and ITT glucose AUC. B. Epididymal fat mass and GTT glucose AUC. C. Body mass and ITT glucose AUC. D. Body mass and GTT glucose AUC. n = 36.

## Discussion

The major finding of our study was that although the HFD induced insulin resistance it did not negatively affect any of the measured mitochondrial characteristics. Rather, markers of both mitochondrial content and respiratory function were increased in HFD animals in both the soleus and gastrocnemius. β-Had activity was also increased in the soleus of HFD animals, but not the gastrocnemius. Although exercise training improved glucose and insulin tolerance in HFD animals, this was not associated with further increases in mitochondrial respiratory function or content. The lack of an association between changes in mitochondrial characteristics and insulin resistance/glucose tolerance was further reinforced by the absence of consistent or strong correlations between these measures.

As cross-sectional studies have reported insulin resistance to associate with decreased mitochondrial content [1–3, 35], we first investigated if HFD-induced insulin resistance was accompanied by a decrease in markers of mitochondrial content. We observed increases in mitochondrial subunit protein expression, as well as CS activity, in the gastrocnemius, indicating an increase in mitochondrial content with HFD. In the soleus, mitochondrial subunit expression was increased by HFD, but CS activity was unchanged. Previous studies have reported both increases [14, 15, 36] and decreases [12] in mitochondrial content with HFD in rodents. Methodological factors such as muscle type examined and the duration and content of the HFD may have contributed to these contrasting findings. Indeed, in this study we found that CS activity was increased in the gastrocnemius muscle, but unchanged in the soleus muscle, which is likely due to differences in metabolism and mitochondrial content between these muscles. It is unclear why patients with insulin resistance and type 2 diabetes display a decrease in muscle mitochondrial content whilst HFD rodents with insulin resistance display an increase. Two main theories prevail. Firstly, that the decrease in muscle mitochondria seen in humans is actually due to physical inactivity [7] or, secondly, that prolonged insulin resistance itself causes a decrease in mitochondrial content [37]. The development of type 2 diabetes is also known to be multi-factorial, and a HFD in rodents only models some aspects. Therefore, further studies in humans are required to tease out the relationship between physical activity, mitochondrial content, and the development of insulin resistance.

As changes in mitochondrial content and function can be dissociated [17, 38], we also investigated the effects of HFD on mitochondrial respiratory function. Mitochondrial respiratory function was increased in both the soleus and gastrocnemius with HFD. This is in contrast to our hypothesis, but some studies have also reported an increase in mitochondrial function with HFD [14, 15, 39]. While two studies have observed a decrease in mitochondrial respiratory function with HFD [11, 12]. This can possibly be attributed to differences in substrates used in the respiration measurements or other methodological differences discussed above (e.g. muscle type examined, type and duration of the HFD). The increase in mitochondrial respiration with HFD is also in contrast to the lower mitochondrial respiration observed in type 2 diabetes patients [5, 6]. However, as discussed in relation to mitochondrial content, the apparently contrasting findings may be due to differences between the HFD model in rodents and type 2 diabetes in human patients.

A number of studies have proposed that chronically elevated fatty acids, intermediates, and increased intramyocellular fat, may play a role in the development of insulin resistance [40]. This is potentially due to either impaired fat oxidation and/or uptake of fatty acids in excess of energy requirements [40, 41]. However, in the current study β-HAD activity was increased with HFD in the soleus, although not the gastrocnemius. Fatty acid oxidation and markers of fatty acid oxidation, such as β-HAD, have also been shown to be increased with HFD in other studies [14, 15]. Thus, it appears in a HFD rodent model of insulin resistance that skeletal muscle mitochondria increase their capacity for fatty acid oxidation in an effort to compensate for the increased supply. This suggests that a decrease in fat oxidation is not responsible for insulin resistance, at least in this HFD model.

Exercise training has previously been reported to reverse impairments in insulin sensitivity/glucose tolerance induced by HFD [23, 25]. This is consistent with our results, where we observed an improvement in both insulin and glucose tolerance in the HFD animals who exercised. However, a novel aspect of our study was to investigate if improvements in insulin and glucose tolerance following exercise training in HFD rats were accompanied by changes in mitochondrial characteristics. We found that eight weeks of exercise training increased mitochondrial respiratory function in the soleus, but not white gastrocnemius in chow fed rats. The differing response between these two muscles may have been due to less recruitment of the white part of the gastrocnemius muscle during the exercise training employed in this study or changes that were below our detectable power. Exercise training did not increase mitochondrial respiratory function in either muscle in the HFD rats, compared to the HFD sedentary rats. Together with no training-induced increase in mitochondrial content in either the soleus or the gastrocnemius muscle, these results suggest that improvements seen in insulin and glucose tolerance are not the result of increases in mitochondrial content or function in HFD rodents.

The lack of an association between changes in mitochondrial characteristics and insulin and glucose tolerance was further reinforced by the absence of consistent or strong correlations between these measures. Previous studies examining correlations between these measures in human participants have found correlations between mitochondrial content and insulin resistance [3], but correlations between mitochondrial respiration and insulin resistance have been less consistent [6] [42, 43]. Body mass and epididymal fat mass did correlate with GTT AUC (i.e. glucose tolerance), indicating that visceral fat mass may be more strongly associated with insulin resistance than mitochondrial characteristics. Visceral fat mass has been previously shown to be an associated with insulin resistance [44].

In this study, we have measured a range of mitochondrial characteristics - none of which were reduced with HFD-induced insulin resistance. While this suggests a decrease in mitochondrial content and respiratory function is not responsible for insulin resistance, it is possible that other mitochondrial characteristics not measured were changed and may contribute to HFD-induced insulin resistance. For example, previous work by Miotto et al [11] in HFD mice has shown that although maximal mitochondrial respiration is increased under high or saturating ADP levels (as in the current study) respiration is actually decreased when lower more biologically relevant ADP levels are used in respiration [11]. We also did not measure mitochondrial reactive oxygen species production, which has been reported to be increased in HFD rodents [45] and has been implicated in the development of insulin resistance [46]. Nor did we examine mitophagy or mitochondrial fission and fusion, which can act as a quality control mechanism for the mitochondria and may cause changes in the efficiency of the mitochondria [47, 48]. However, what is clear is that the upregulation of mitochondrial content and respiratory function we observed in skeletal muscle in HFD rodents is most likely a compensatory response to the increased lipid availability, but which may not be sufficient to prevent ectopic lipid accumulation and insulin resistance.

In conclusion, while findings from cross-sectional studies have indicated a potential link between altered mitochondrial characteristics and insulin resistance, our results do not provide support for a causal link between HFD-induced insulin resistance and altered mitochondrial characteristics. In contrast, HFD-induced insulin resistance was associated with increased mitochondrial content, β- HAD activity, and mitochondrial respiration. Furthermore, exercise training-induced improvements in insulin resistance were not accompanied by parallel improvements in mitochondrial characteristics. Our results suggest that decreases in mitochondrial content and function do not precede and are not necessary for the development of skeletal muscle insulin resistance induced by HFD in rodents.

## Nonstandard abbreviations

β-HAD: Beta(3)-hydroxyacyl-CoA dehydrogenase
CS: citrate synthase
CI: complex I
CI+II: convergent electron input through CI and CII
E: electron transport system (ETS) capacity
L: leak respiration
P: oxidative phosphorylation capacity
mtDNA: mitochondrial DNA
TBS: tris buffered saline
TEM: technical error of measurement

## Acknowledgements

Thank you to Jarrod Kerris, Kelly Wilson, and visiting students for their assistance with animal work.

## Data availability

The data that supports the findings of this study is included within the article and/or available from the corresponding author upon reasonable request.

## Author contributions

A.G. contributed to the study conception and design, data collection, and interpretation and analysis and drafting of the manuscript. J.K, E.M., N.S., J.B., M.J., V.A-S., and J.C. contributed to data collection. G.M. contributed to study design and interpretation. D.B. contributed to study conception, design, and interpretation and drafting of the manuscript. All authors critically revised the manuscript and approved the final version.

